# An Artificial Intelligence and Telemedicine Based Screening Tool to Identify Glaucoma Suspects from Color Fundus Imaging

**DOI:** 10.1101/2021.04.16.440184

**Authors:** Alauddin Bhuiyan, Arun Govindaiah, R Theodore Smith

## Abstract

**Backgrounds & Objective:** Glaucomatous vision loss may be preceded by an enlargement of the cup-to-disc ratio (CDR). We propose to develop and validate an artificial intelligence based CDR grading system that may aid in effective glaucoma-suspect screening.

**Design, Setting & Participants:** 1546 disc-centered fundus images were selected including all 457 images from the Retinal Image Database for Optic Nerve Evaluation dataset, and images randomly selected from the Age-Related Eye Disease Study, and Singapore Malay Eye Study to develop the system. First, a proprietary semi-automated software was used by an expert grader to quantify vertical CDR. Then, using CDR below 0.5 (not suspect) and CDR above 0.5 (glaucoma-suspect), deep learning architectures were used to train and test a binary classifier system.

**Measurements:** The binary classifier, with glaucoma-suspect as positive, is measured using sensitivity, specificity, accuracy, and AUC.

**Results:** The system achieved an accuracy of 89.67% (sensitivity, 83.33%; specificity, 93.89%; AUC, 0.93). For external validation, the Retinal Fundus Image database for Glaucoma Analysis dataset, which has 638 gradable quality images, was used. Here the model achieved an accuracy of 83.54% (sensitivity, 80.11%; specificity, 84.96%; AUC, 0.85).

**Conclusions:** Having demonstrated an accurate and fully automated glaucoma-suspect screening system that can be deployed on telemedicine platforms, we plan prospective trials to determine the feasibility of the system in primary care settings.

## 1. Introduction

Glaucoma is a group of diseases that damage the eye’s optic nerve and can result in vision loss and blindness [1]. Glaucoma, with age-related macular degeneration (AMD) and diabetic retinopathy (DR), is one of the three leading causes of blindness in developed countries, and is now the second leading cause of blindness globally, after cataracts [2, 3].

Glaucoma is characterized by loss of retinal ganglion cells (RGCs), which results in visual field impairment and structural changes to the retinal nerve fiber layer (RNFL) and optic disc [4]. Glaucoma has few early symptoms; Over 3 million Americans have glaucoma, and the number is over 76 million worldwide, with projections showing 111 million by 2040 [5]. About half of those affected do not know it [6]. Most of the time, when detected, it is already late, i.e., with irreversible visual field loss. Therefore, it is essential to identify individuals at the early stages of this disease for treatment. The social and economic costs of vision loss from glaucoma are also extremely high. Early detection of these conditions halts a downward spiral in overall health: depression, loss of independence, need for nursing home care, falls, fractures, and death. These adverse outcomes are also extremely costly. The total economic burden in the US, direct and indirect, of vision loss and blindness from all causes, is now $145 billion, expected to triple by 2050 in real dollars, with increased longevity generally [7].

The relationship between estimated RGC counts and CDR suggests that assessment of change in CDR is a sensitive method for the evaluation of progressive neural losses in glaucoma, specifically the retinal cup-disc ratio (CDR) is highly correlated with glaucoma [8-13]. Although several techniques [8, 9, 12-14] have been proposed to measure the cup-disc ratio, they have not been extensively validated for screening, and the current research is mainly focused on overall glaucoma detection. For example, Saxena et al. [15] proposed a method for glaucoma detection using the standard ORIGA dataset in 2020, whose AUC is 0.82. However, the American Academy of Ophthalmology outlined a set of tests to determine glaucoma which are: eye pressure, eye drainage angle, optic nerve damage, peripheral side vision (visual field test), computerized imaging of optic nerve, and thickness of the cornea [16]. Thus, using color fundus imaging alone is not a standard protocol for glaucoma detection or diagnosis. The CDR can be an effective tool to identify the glaucoma suspect and our focus is mainly to identify the glaucoma suspect individuals (as a screening process from primary care settings), who can be further tested to determine glaucoma and its progression. Thus, starting with cup-to-disc ratio may also be more suited for clinical applicability, because of its inherent explicability. At present, although there are techniques to detect CDRs, a full-fledged low-cost automated system that passively detects glaucoma from patients’ yearly visits in a primary care setting is still not widely available

We have considered the vertical cup-disc ratio and the threshold value to determine the normal and abnormal, based on the following studies. A larger or abnormal CDR is mentioned in [10] and categorized as CDR>0.5. The same research scheme also suggested that small changes in CDR may be associated with significant losses of RGCs, especially in eyes with large CDRs. Enlarged CDR is one indicator of the risk of glaucoma [11]. Most individuals fall near the average vertical CDR of 0.4, and 2.5% of the population have a cup/disc ratio of over 0.7 [17]. Studies showed that for the normal (non-glaucoma) population, the horizontal C/D ratio is usually larger than the vertical C/D ratio, but the vertical ratio increases faster in early and intermediate stages of glaucoma [21]. Also, studies have documented that the normal C/D ratios ranging from less than 0.3 (66 percent of normal individuals) to greater than 0.5 (only 6 percent of normal individuals). Therefore, we considered CDR 0.5 and above as glaucoma suspect [22, 23] and began exploring deep learning methods for detecting larger CDRs [24]

We note that we have utilized the quantified vertical CDR when other research schemes used the qualitative assessment (e.g., small, medium, and large). We developed and validated our vertical CDR quantification software to perform this quantified grading. The software demonstrates high repeatability and reliability, which we have also provided in the paper. We developed and validated our AI-based glaucoma suspect screening results based on the quantified vertical CDR. This should provide higher accuracy and confidence than selective judgment.

- The paper describes a method for glaucoma suspect screening which utilizes a cloud-based system and incorporates the telemedicine facilities. Thus, the screening will be available in remote clinics and primary care settings.
- The paper describes results on a novel automated method that addresses the early screening of glaucoma suspects which is a major public health concern.
- Therefore, an accurate and efficient screening in the remote primary care settings can provide a mass screening of the population who are currently dropping from yearly visits to the ophthalmologist.

The rest of the paper describes the development and validation of this glaucoma suspect screening tool.

## 2. Materials and Methods

### The global strategy of the study is organized as follows

2.1 Data sources, describing the various datasets,

2.2 Ground truth, describing manual grading

2.3 Preprocessing, describing data curation and data processing before training, and

2.4 Architecture, describing the technical details of the training and validation.

#### 2.1. Data Sources

Fundus images from three sources were used to conduct training experiments and a fourth for external validation. A total of 1546 color fundus images that included the disc were selected randomly from the Age-Related Eye Disease Study (AREDS) [37] and Singapore Malay Eye Study (SiMES) study [38], and all the images from Retinal Image Database for Optic Nerve Evaluation (RIM-ONE) dataset [39], an ophthalmic reference image database specifically designed for glaucoma analysis. For external validation, we used the Online Retinal fundus Image database for glaucoma analysis (ORIGA) [40]. Although these retinal images had already been graded for glaucoma, we performed our gradings for consistency (Section 2.2).

Briefly, AREDS is a 13-year study of age-related eye diseases. The participants were of the ages 55 to 80 when they were enrolled. 30-degree fundus photographs were graded as glaucoma present or absent by the AREDS ophthalmic grading center. We used fundus images from those cases as well as from the normal control population for this experiment.

SiMES-1 was a cross-sectional, population-based epidemiological study of eye diseases. It was performed on 3,280 randomly selected Malay adults living in the south-western part of Singapore. All study participants underwent various questionnaires and detailed eye examinations. We have taken those images for which information about the presence or absence of glaucoma was present.

RIM-ONE is an ophthalmic reference image database specifically designed for glaucoma diagnosis, not only for medical education purposes but also as an evaluation tool for designers of segmentation algorithms. RIM-ONE is available as a free download as part of a research collaboration between three Spanish hospitals: Hospital Universitario de Canarias, Hospital Clínico San Carlos, and Hospital Universitario Miguel Servet.

The ORIGA-light dataset is an ophthalmic reference image database specifically designed for glaucoma analysis. ORIGA-light serves as a benchmarking resource for researchers to evaluate image processing algorithms that detect and analyze various image signs highly related to glaucoma diagnosis. To facilitate this, the authors of ORIGA used their in-house grading tools to grade several glaucoma-related signs. The publicly available dataset that we used has 680 graded images, out of which 460 are healthy and the rest are graded as glaucoma, taken from adults aged between 40 and 80 years. Each image is segmented and annotated by trained professionals from the Singapore Eye Research Institute.

AREDS dataset may be obtained with request on their website – “dbgap.ncbi.nlm.nih.gov”. All the other datasets are available upon request from the authors of the corresponding datasets.

#### 2.2. Ground truth

As noted, we did not use prior annotations of the presence of glaucoma but instead graded each image manually for vertical and horizontal CDR. A proprietary software called “CDR annotator” [36] was used for the purpose. Figure 1 shows the interface for marking the region of the cup and the disc, from which the vertical and horizontal CDRs are automatically generated. Before this, regions of interest (optic disc) were identified and cropped from the fundus image automatically using custom deep learning methods (unpublished, Figure 2).

**Figure 1.**
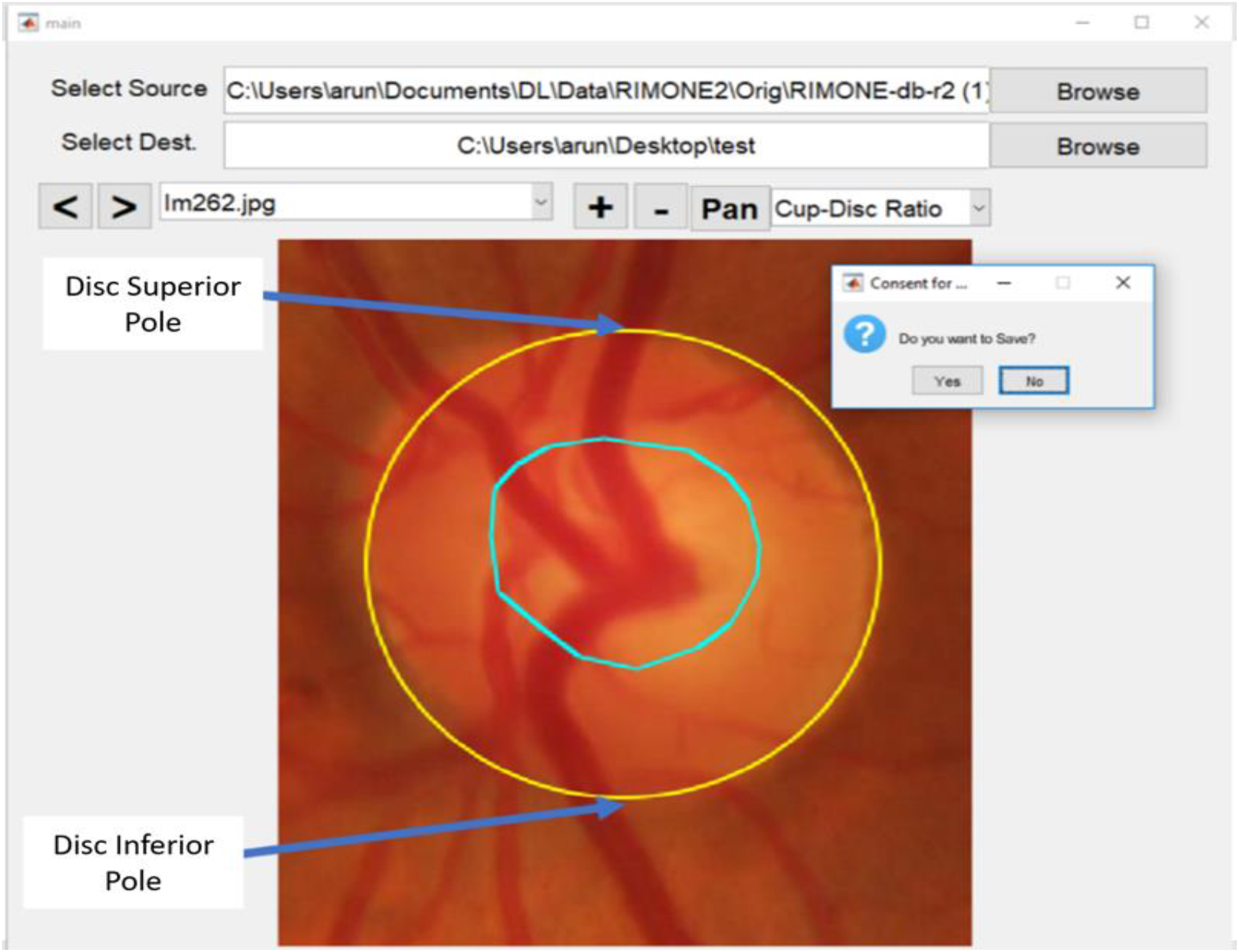
The interface of “CDR annotator”, the proprietary cup/disc ratio grading software. The optic cup is circled in blue.

**Figure 2.**
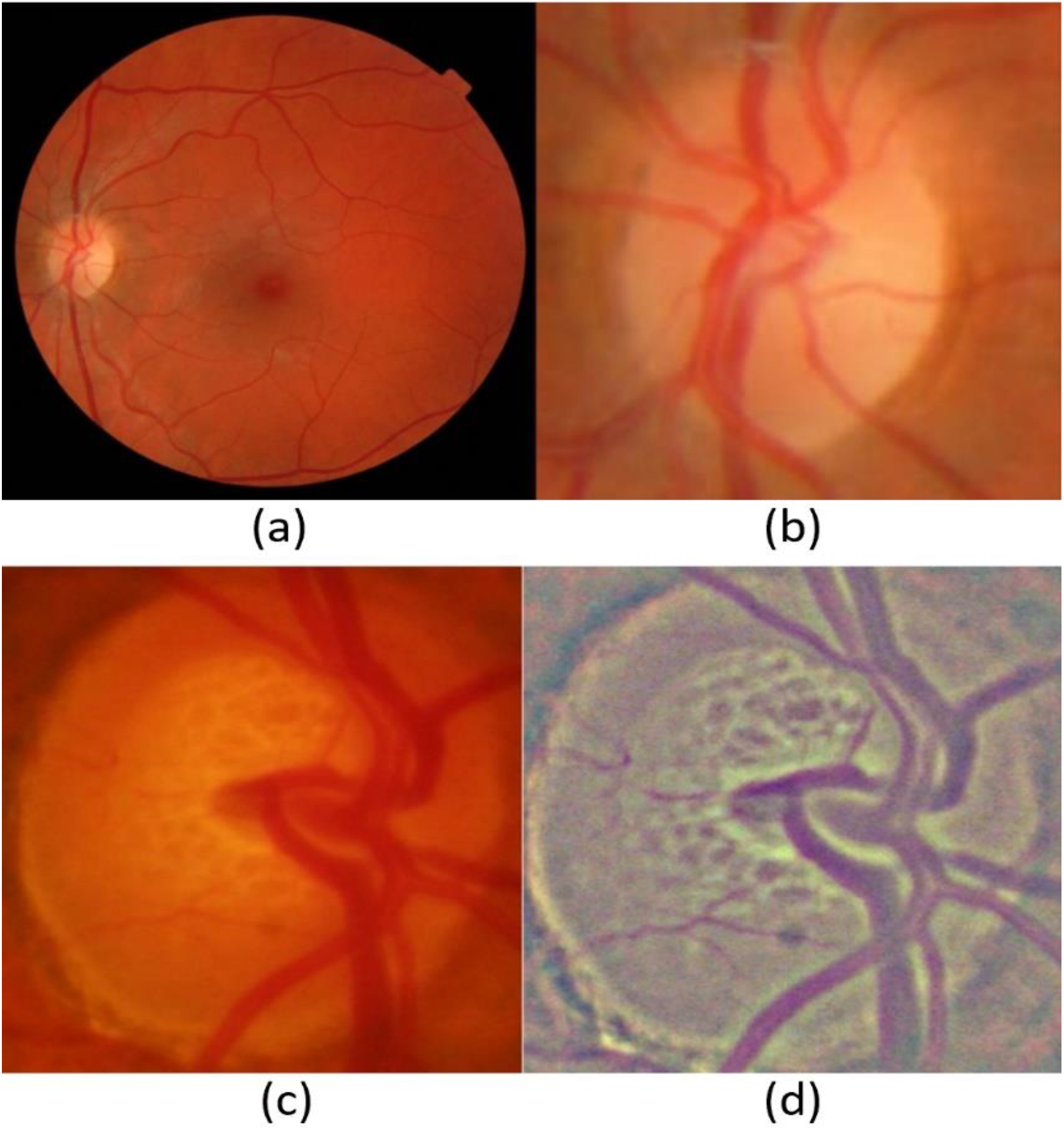
A 30-degree fundus image (a) and the automatically cropped optic disc region (b). (c) shows another cropped example that is preprocessed to obtain the final image (d).

Two computer engineers at iHealthScreen Inc. were trained by expert ophthalmologists to grade CDRs in each image. Whenever there was disagreement in grading, the two graders adjudicated and produced uniform grading (CDRs) as ground truth for the images. Before adjudication, 250 images were randomly chosen to evaluate intergrader correlation and 200 for intra-grader correlation. The intergrader and intra-grader Pearson correlations between their CDR ratio annotations, 0.832 and 0.841 respectively [41], showed good consistency. The vertical CDRs were used to categorize the images in two classes: Class 1 (not glaucoma suspect): vertical CDR ≤ 0.5 and class 2 (glaucoma suspect): vertical CDR > 0.5. After the categorization, the final dataset used for training was as shown in Table 1. Similarly, after quality control (removing 42 ungradable images), 638 images were selected out of a total of 680 images in the ORIGA-light dataset and processed and graded. CDR’s of 452 images in the ORIGA dataset were graded by the experts to be less than or equal to 0.5, and those of 186 images were graded to be above 0.5.

**Table 1.** Number of images taken from AREDS, SIMES, and RIMONE1. The table shows the various groups to which the images belong based on their graded cup/disc ratios.

| Category (manually graded for vertical<br>cup/disc ratio) | AREDS | SIMES | RIMONE1 | Total |
| --- | --- | --- | --- | --- |
| $\leq 0.5$ | 462 | 253 | 342 | 1057 |
| above 0.5 | 131 | 243 | 115 | 489 |

#### 2.3. Preprocessing

Two types of input images were used simultaneously –the original RGB and a transformed, or preprocessed, RGB image. The transformed RGB image is a color space averaged image [42]. We subtracted the local means with a kernel size 8 and Gaussian blurring. Such a preprocessing technique of local color averages (LCA) is effective when dealing with images from various sources taken under different conditions. Figure 2 (c & d) shows an example of such a preprocessing technique.

The images from three datasets were combined to form a unified dataset. The test-set was then taken from these images randomly. These cropped images from the original RGB image set and the LCA set form the final input for this experiment.

#### 2.4. The architecture and the telemedicine platform

The architecture we propose (shown in Figure 3) consists of an ensemble of five neural networks and a classical tree learning algorithm. The overall system is a binary classifier that classifies the images into one of the two categories (CDR <= 0.5 and CDR>0.5).

**Figure 3.**
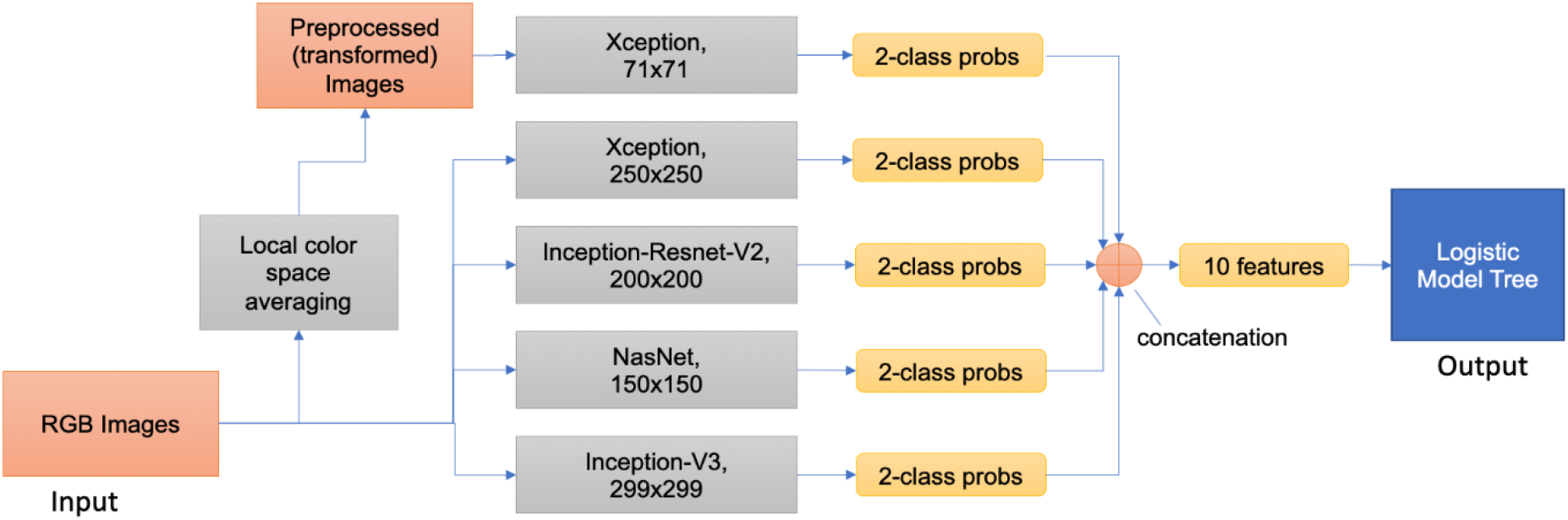
Overall schematic representation of the model building. The figure shows the two types of input images (RGB and transformed) fed into five neural networks. Each network accepts a rescaled image (shown below the name of the architecture) and produces two probabilities. The ten resulting probabilities are then concatenated to form a feature array of size 10 and fed as input to a logistic model tree which acts as the final classifier in this system.

To build an image classification model robust in terms of image and dataset variations, and that is capable of learning features on such a wide scale in terms of size and location, the image preprocessing techniques and neural network selections were made carefully [43]. Multiple different neural networks, when used, are hypothesized to learn features from an image differently. Combining the results from different models to produce a final output is a general practice to obtain a better performance than each of the constituent network architectures [44]. To increase the robustness, different input sizes for the networks were chosen. Also, two types of images are fed into the models. One type is regular RGB images, and the other is preprocessed LCA images.

Deep learning architectures were trained and validated to produce completely automatic outputs. All graded images (1546) were grouped into two categories: CDR ≤ 0.5 (1057 images) and CDR above 0.5 (489 images) in the two-class model.

The initial test models (from which the final models are chosen) were built to evaluate feasibility. The network architectures used are Inception-Resnet V2 [45], NasNet [46], Xception [47] and Inception [48]. Deep learning architectures like alexnet [49] and VGG networks [50] initially focused on stacking layers deeper and deeper, hoping to get better performance. Inception architecture changed this approach by going “wider” instead of focusing on going just “deeper”. Every version of inception optimized speed and accuracy and the version used in our experiment – Inception V3 – uses RMSProp optimizer [51], factorized 7 × 7 convolutions, and added batch normalization [52] in the auxiliary classifiers.

The original Inception networks without depth-wise separable convolutions were modified to include the residual connections, called the Inception-ResNet networks. Inception modules allowed for adding more layers, and residual connections made the network converge faster, which is the basis for Inception-Resnet V2. Xception is a novel deep convolutional neural network architecture inspired by Inception, but where Inception modules have been replaced with depth-wise separable convolutions. This architecture slightly outperforms Inception V3 on the ImageNet dataset. NASNet learns the model architectures directly on the dataset of interest by searching the best convolutional layer (or “cell”) on a small dataset and then applying it to a larger dataset by stacking together more copies of this cell.[53]

The various input sizes used range from 71×71 to 399×399. The best models thus obtained will be ensembled to form a final architecture. Through experimentation, we developed an ensemble of five networks in the final architecture for the glaucoma screening system. The description of the five models is given below.

1. Xception, Input size: 71×71, Input Image Type: Local Color Averaged (transformed)
2. Xception, Input size: 250×250, Input Image Type: RGB
3. Inception-Resnet-V2, Input size: 200×200, Input Image Type: RGB
4. NasNet, Input Size: 150×150, Input Image Type: RGB
5. Inception-V3, Input size: 299×299, Input image Type: RGB

The full framework for building the model is shown in Figure 3. The networks are trained for 500 epochs. We trained the networks with a batch size of 20 images. This is a high number considering the limitations of GPU memory. The low resolution of cropped images helped achieve a bigger batch size. The Adam [54] optimizer was used with a learning rate of 0.0004. To save time, an early stopping mechanism halted training if there was no improvement for 20 consecutive epochs. Every epoch is monitored for loss and this value is used for early stopping. The loss function used in this system was categorical cross-entropy. This quantity was used to determine the hyperparameters of the networks. SoftMax activation is used as the last layer in each of the architectures. All the networks were trained on NVIDIA Titan V GPU for two weeks with an average time of 20 to 30 minutes per epoch.

Each model gives a probability array of size 2 for 2 classes. The five arrays from five models are concatenated to form a feature array of size 10 that is then used to build a Logistic Model Tree [55] model for final output. Sensitivity, specificity, accuracy, and Cohen’s kappa were calculated to evaluate the models.

#### 2.5 The role of the AI platform

The CDRcarries three advantages: first, it is a single variable that is known to be strongly correlated with the disease, in particular with losses of RGCs as noted, and as such, is inherently explicable and acceptable to the eye community; second, measurement of CDR can be accomplished from a single retinal color photograph obtained by an automated, non-mydriatic camera in a primary care office and forwarded on a telemedicine platform for expert interpretation with semi-automated methods [36]; third, that expert interpretation, which is still time-consuming and expensive for humans, can be replaced by AI for efficiently and accurately evaluating the images as we propose to demonstrate herein. In fact, we have already introduced such a HIPAA-compliant telemedicine platform, iPredict, with the requisite capabilities of AI solutions and report generation.

A telemedicine platform has been introduced that enables the cloud-based processing of the AI solutions and report generation can extensively simplify the process of evaluating the images on a mobile/tablet or a low-performance computer, a requirement for the successful glaucoma suspect screening at primary care settings. We aim to address this with our HIPAA compliant telemedicine platform iPredict.

In the future, we propose to use the Software Tool ‘iPredict-glaucoma’ at iPredict platform (https://ipredict.health/): An online version of the Glaucoma Suspect Screening system is available at https://www.ihealthscreen.org/ipredict-glaucoma/ (the username and password can be obtained for research purposes through writing the corresponding author). The AI-based telemedicine platform iPredict developed by iHealthscreen Inc. integrates the server-side programs (the image analysis and deep-learning modules for screening systems) and local remote computer/mobile devices (for collecting and uploading patient data and images). The images are first checked for gradability automatically by an artificial intelligence based system developed in-house from 3000 fundus images manually graded for gradability, the system achieved over 99% accuracy. The server analyzes the images, and a report will be sent to the remote clinic with an individual’s screening results and further recommendations.

## 3. Results

The two-class glaucoma model (CDR ≤ 0.5 and above 0.5) achieved an accuracy of 89.67% (95% CI - 85.65% to 92.87%) with a sensitivity of 83.33% (95% CI - 75.44% to 89.51%) and a specificity of 93.89% (95% CI - 89.33% to 96.91%) along with a Cohen’s kappa of 0.78 (95% CI - 0.71 to 0.85) when above 0.5 cases are considered as positive. AUC for the same data was 0.93 (0.89 to 0.96) as shown in Figure 4. The complete results of the system are detailed in Table 2.

**Figure 4.**
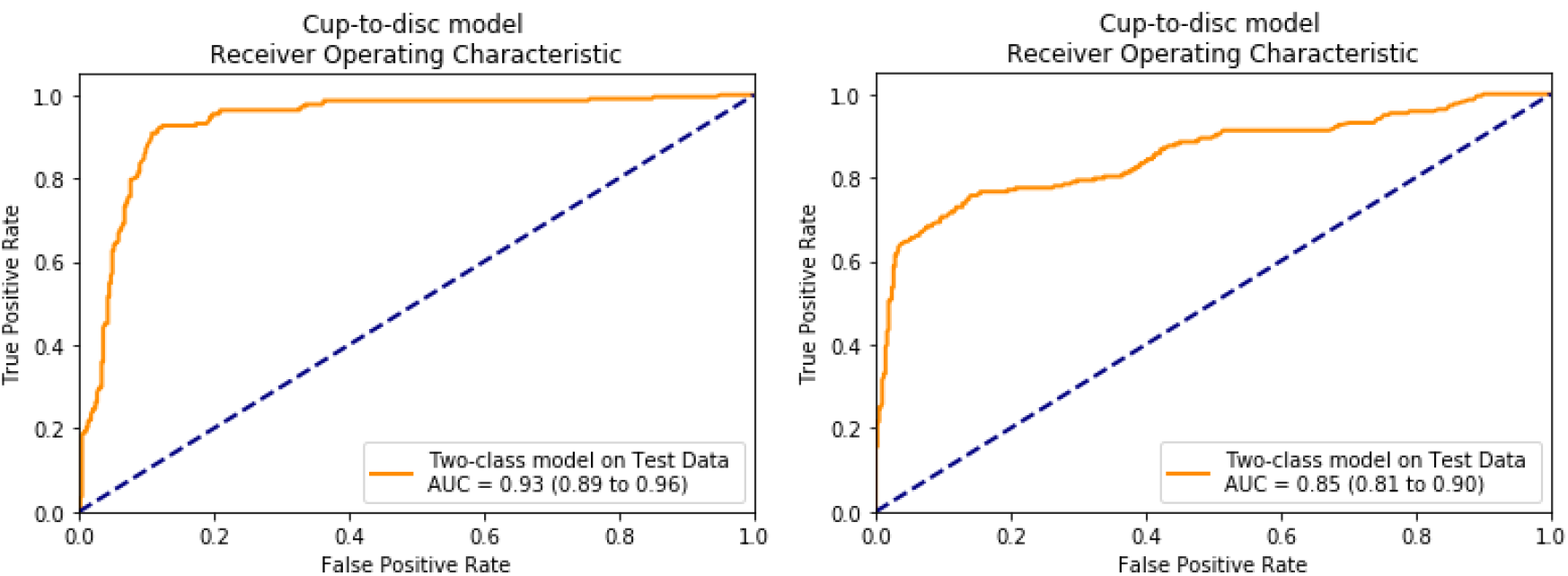
Area under ROC of the test data results (left) and the results for ORIGA-light dataset (right) for the two-class cup-to-disc ratio model.

**Table 2.** Accuracy, sensitivity, specificity, Cohen’s kappa, and AUC for the system with 95% confidence intervals on the test data and ORIGA-light (validation data), with the ratio over 0.5 considered as a positive case for the model.

| Measure | Test data (with 95% CI) | ORIGA-light (with 95% CI) |
| --- | --- | --- |
| Accuracy | 89.67% (85.65% to 92.87%) | 80.11% (73.64% to 85.59%) |
| Sensitivity | 83.33% (75.44% to 89.51%) | 84.96% (81.32% to 88.12%) |
| Specificity | 93.89% (89.33% to 96.91%) | 83.54% (80.43% to 86.34%) |
| Cohen’s kappa | 0.78 (0.71 to 0.85) | 0.62 (0.57 to 0.67) |
| Area Under the Curve | 0.93 (0.89 to 0.96) | 0.85 (0.81 to 0.90) |

On the external validation dataset, the two-class model achieved a sensitivity of 80.11% (73.64% to 85.59%) and specificity of 84.96% (81.32% to 88.12%) with a Cohen’s kappa of 0.62 (0.57 to 0.67) on the ORIGA dataset. AUC for the same data was 0.85 (0.81 to 0.90), as shown in Figure 4. The complete results on the external validation dataset can be found in Table 2.

The cloud-based and HIPAA-compliant telemedicine platform ‘iPredict’ (https://ipredict.health/) has been validated for image and data transfer accuracy. We have transferred and analyzed nearly 850 images for AMD screening and DR screening from 4 primary care clinics in Queens and Manhattan, New York, USA. The initial results from utilizing our platform are reported in [56, 57]. We found a 100% correlation between the results obtained from directly evaluated images and the images transferred and processed by iPredict. We have also tested 100 images for vertical CDR computation and received the same accuracy.

## 4. Discussion

In this study, we have demonstrated an accurate and fully automated deep learning screening system for glaucoma suspects through retinal photography that may be effective for the identification of glaucoma suspects in primary care settings.

Glaucoma is a prevalent, blinding disease worldwide with few symptoms until irreversible later stages and is undiagnosed at rates approaching 50% even in developed countries [6]. Hence the pressing public health need for effective community screening. The need is even greater in communities of color, with an overall ratio of 8:1 for nonwhite to white primary glaucoma blindness, due at least in part to receiving medical care later in the disease than whites [58], Compounding the problem, there is also a dramatically earlier age of onset in this group. In an Afro-Caribbean population glaucoma suspect status was high across all age groups, with significant prevalence even in populations less than 40 years of age [59].

We have shown on several large datasets that the cup/disc ratio (CDR) can be measured automatically from retinal photography with sufficient accuracy to discriminate suspects from non-suspects and thus potentially facilitate referral of suspects to an ophthalmologist for specialized care. Thus, a future, achievable goal is an AI *telemedicine* platform in which our current methodology will be deployed in primary care settings through remote image capture. A prospective trial will be needed to determine the feasibility of the system in clinical settings, with inexpensive, automated non-mydriatic retinal cameras and a telemedicine platform for image transfer to the deep learning screening system. Such systems have been tested clinically with proven accuracy for screening DR in comparison to expert graders [29]. It is thus reasonable to expect that similar success may be achieved with glaucoma.

A National Eye Institute study showed that 90% of glaucoma subjects can be prevented from progression to severe glaucoma through timely identification and intervention [6] However, nearly sixty percent of Americans with diabetes skip annual sight-saving exams recommended by their Primary Care Physicians (PCPs). Given such poor compliance by diabetics, who are informed about the risks to their vision, it is likely that compliance with eye exams is even worse in the general population [60]. Therefore, our focus is to identify the suspects in the primary care settings, not only to get them needed care but also to eliminate large numbers of unnecessary specialist visits for glaucoma screening.

The medical imaging and diagnostics field has been revolutionized by advances in deep learning in recent years. Extensive research interest is being shown in using artificial intelligence for solving medical problems [25]. Ting et al. [26] detailed the potential applications of AI in ophthalmology. Gulshan et al. [27], in their seminal paper, showed the application of AI in diabetic retinopathy from fundus images using deep learning. Recently, we have published two groundbreaking works on late AMD prediction [28] and diabetes screening in primary care settings [29]. There is also considerable research in other medical areas such as multiple sclerosis [30], neurodegeneration [31], and age-related macular degeneration [32-35]. Several AI techniques [8, 9, 12-14] have been proposed to measure the cup-disc ratio, ; they have not been validated for screening glaucoma suspects.

We note that the current research using color fundus imaging is mainly focused on the detection of glaucoma, which we believe is not an appropriate option for clinical settings if we follow the glaucoma detection or diagnosis protocol (https://www.aao.org/eye-health/diseases/glaucoma-diagnosis). Thus, here we aim to clear the differentiation of the term glaucoma and glaucoma suspect. The glaucoma detection is a diagnosis of glaucoma that requires the structural and functional abnormalities from glaucoma suspect, which is implied by its name, a category of markers with an increased likelihood of the disease.

Li et al. [18] and Ting et al. [19] trained computer algorithms to detect the glaucoma-like disc, defined as a vertical CDR of 0.7 and 0.8, respectively. In general, an eye with vertical CDR above 0.5 is considered a glaucoma suspect [20]. In this paper, we introduced an automated cup-disc measurement tool which can determine if the vertical cup-disc ratio is above or below 0.5, in conjunction with a deep machine learning based tool. We have published this approach, an ensemble of deep learning architectures with a logistic tree at the end, for effective use in AMD screening, but it is novel in glaucoma screening. With a telemedicine platform, it could provide screening of glaucoma suspects on a large scale in primary care settings, conferring substantial public health benefits of reduced vision loss from glaucoma and reduced health care costs.

### Limitations and Future Work

In our testing, the sensitivities are somewhat lower than the specificities, with therefore a somewhat greater risk of missing true cases. In our future work in this project, we aim to understand and anticipate the doctors’ requirements in this new method of screening and tune the system such that the false positives and false negatives are in an acceptable ratio. Small discs with “pathologic cups” are hard to detect. Geometrically, a small disc with a CDR of 0.7 has much less healthy neural rim tissue than a normal-sized disc with the same CDR. Therefore, CDR asymmetry would be a reasonable addition to the screening program. In general, a provisional diagnosis of glaucoma suspect is generally given with CDR asymmetry (>/=0.20) [61]. This criterion could be implemented in the next version of the present DL architecture that is already tuned to CDR measures. Asymmetrical cupping of the optic disc was found in 5.6% of normal individuals, in 30% of a group with ocular hypertension without field defect, and in 36% of those with established chronic open-angle glaucoma and field loss [62]. We note that our algorithm classifies an optic disc based on vertical CDR based on single retinal image, and asymmetrical cupping may show up as different reading in the two eyes of the same image - in turn, helping doctors with an additional biomarker for glaucoma.

### Strengths

The model was built on several large datasets, with external validation on another. The output is an ideal binary target for glaucoma suspects with a single highly correlated and easily measured/interpreted variable. We have had success with a novel hybrid AI approach, to screening for AMD and DR from fundus photos, that performs at least as well as other techniques in the literature, and so we chose this route again for screening glaucoma suspects: an ensemble of DL techniques is first trained on the image inputs to produce sets of probabilities (one set for each DL technique) for classifying the image into a disease state; these sets of probabilities are then inputted to an independently trained logistic model tree which acts as the final classifier in this system. Aglaucoma diagnosis requires multiple structural and functional criteria that are not available or appropriate in the primary care setting. To our knowledge, our system is the only one proposed that is full-fledged, passive screening system for an adequate screening of glaucoma suspects with a single disease marker that could be easily obtained in the primary care setting on a telemedicine platform without expensive, specialized equipment or services.

Future work can carry these methods into the primary care setting to perform annual screening for this silent blinding disease. We propose to address this urgent public health need with future prospective trials of our system for low-cost, rapid glaucoma screening. These trials will be modeled on our current ongoing NIH funded 3-year trial (SBIR Phase IIb R44EY031202, A Bhuiyan PI, “https://projectreporter.nih.gov/project_info_details.cfm?aid=10010769") for detection and prediction of AMD in primary care settings with our published DL algorithms [28]. A complete HIPAA compliant functional AI-based telemedicine platform for real-time diagnosis is already in place which integrates the server-side screening programs (image analysis and deep-learning modules) and local remote devices (for collecting patient data and images).

## 5. Conclusions

We have developed an effective deep learning/logistic model tree hybrid screening tool for the identification of glaucoma suspects by vertical CDR from non-mydriatic retinal photographs.

Building on this tool, a full AI telemedicine platform is envisioned in a future state in which our current AI methodology will be deployed in primary care settings. Fully automated cameras will capture images for transfer through the cloud to the server-side for immediate results and further patient referral if needed, with significant public health benefit for early detection and prevention of this sight-threatening disease.

## 6. Acknowledgements

## Author Contributions

Conceptualization, A.B., and A.G.; methodology, A.B., and A.G.; validation, A.G., and A.B.; formal analysis, A.G.; investigation, A.G.; data curation, A.B. and R.T.S.; writing— review and editing, A.G., R.T.S., and A.B.; supervision, R.T.S.; project administration, A.B.; funding acquisition, A.B. All authors have read and agreed to the published version of the manuscript.

## 7. Data availability

The data that support the findings of this study are available from the corresponding author upon reasonable request.

